# scMUSCL: Multi-Source Transfer Learning for Clustering scRNA-seq Data

**DOI:** 10.1101/2024.04.22.590645

**Authors:** Arash Khoeini, Funda Sar, Yen-Yi Lin, Colin Collins, Martin Ester

## Abstract

**Motivation:** scRNA-seq analysis relies heavily on single-cell clustering to perform many downstream functions. Several machine learning methods have been proposed to improve the clustering of single cells, yet most of these methods are fully unsupervised and ignore the wealth of publicly available annotated datasets from single-cell experiments. Cells are high-dimensional entities, and unsupervised clustering might find clusters without biological meaning. Exploiting relevant annotated scRNA-seq dataset as the learning reference can provide an algorithm with the knowledge that guides it to better estimate the number of clusters and find meaningful clusters in the target dataset.

**Results:** In this paper, we propose Single Cell MUlti-Source CLustering, scMUSCL, a novel transfer learning method for finding clusters of cells in a target dataset by transferring knowledge from multiple annotated source (reference) datasets. scMUSCL relies on a deep neural network to extract domain and batch invariant cell representations, and it effectively addresses discrepancies across multiple source datasets and between source and target datasets in the new representation space. Unlike existing methods, scMUSCL does not need to know the number of clusters in the target dataset in advance and it does not require batch correction between source and target datasets. We conduct extensive experiments using 20 real-life datasets and show that scMUSCL outperforms the existing unsupervised and transfer-learning-based methods in almost all experiments. In particular, we show that scMUSCL outperforms the state-of-the-art transfer-learning-based scRNA-seq clustering method, MARS, by a large margin.

**Availability:** The Python implementation of scMUSCL is available at https://github.com/arashkhoeini/scMUSCL

## Introduction

Sequence analysis plays a fundamental role in molecular biology since both the genome and proteins have an underlying sequential structure. Tasks of sequence analysis are diverse, including the detection of various types of mutations and the measurement of gene expression levels. The expression level of a gene is given by the number of transcripts of the gene, which may be translated into the corresponding proteins, and provides valuable information about various biological processes and cellular functions. Traditional technologies perform sequence analysis of samples, and return aggregate measurements, e.g., the average expression level of a gene across the cells of a sample, where the transcriptional heterogeneity of the samples are lost. Recently, technologies for sequence analysis of individual cells have emerged, which perform sequence analysis at a much finer level of resolution of individual cells (Eberwine *et al*., 2014). In particular, single-cell RNAseq (scRNA-seq) is a technology that returns the gene expression profiles of individual cells in a given sample.

While scRNA-seq data provide information at a great level of resolution, it comes with the following two challenges. First, scRNA-seq data is very high-dimensional, since the number of genes is in the order of tens of thousands, i.e. roughly 30,000 for humans. Consequently, methods for the analysis of scRNA-seq data typically perform dimensionality reduction. Second, scRNA-seq data is very noisy. Several factors contributes to the noise, including the stochasticity of the transcription, technical factors, such as mRNA capture rate and low accuracy in measuring genes that are expressed only at a low level. Clustering cells alleviates the effect of the noise and is an essential step before downstream tasks, i.e., cell-type annotation, visualization, trajectory analysis and determination of transcriptional heterogeneity in health and disease (Luecken and Theis, 2019; Clarke *et al*., 2021; Wang *et al*., 2019).

Clustering is an unsupervised problem of finding groups such that members of a group are more similar to the other members of the same group than to the members of other groups. Multiple methods have been introduced in the literature to find clusters of cells, or to find a low-dimensional representation which facilitates clustering (Wang *et al*., 2018; Tian *et al*., 2019; Wan *et al*., 2023; Ciortan and Defrance, 2021). One important caveat common to all these methods is that they are fully unsupervised and ignore the wealth of information in publicly available annotated scRNA-seq datasets (Grabski *et al*., 2023; Zhang *et al*., 2023). Exploiting available annotated data can guide a clustering algorithm to find more accurate and biologically meaningful clusters (e.g., clusters that correspond to certain cell-types). However, it is not trivial how to incorporate existing annotated scRNA-seq data into a clustering algorithm. One major problem with leveraging available scRNA-seq datasets is the existing discrepancies between datasets coming from different species, different tissues of the same species, or different sequencing technologies or laboratories. For example, MARS (Brbić *et al*., 2020) aims to utilize multiple annotated scRNA-seq datasets as source domains to learn a clustering for an unannotated target domain. However, MARS does not explicitly align source datasets with each other and with the target dataset, and it suffers from distribution discrepancies (e.g., batch effect) in the source and target datasets (illustrated in Figure 1a). This limits MARS’s ability to learn from common source clusters and to distinguish them from new clusters that appear only in the target domain. Further, almost all previous works on cell clustering presume the true number of clusters in advance, which is a major drawback since in practice the actual number of clusters is unknown.

**Fig. 1.**
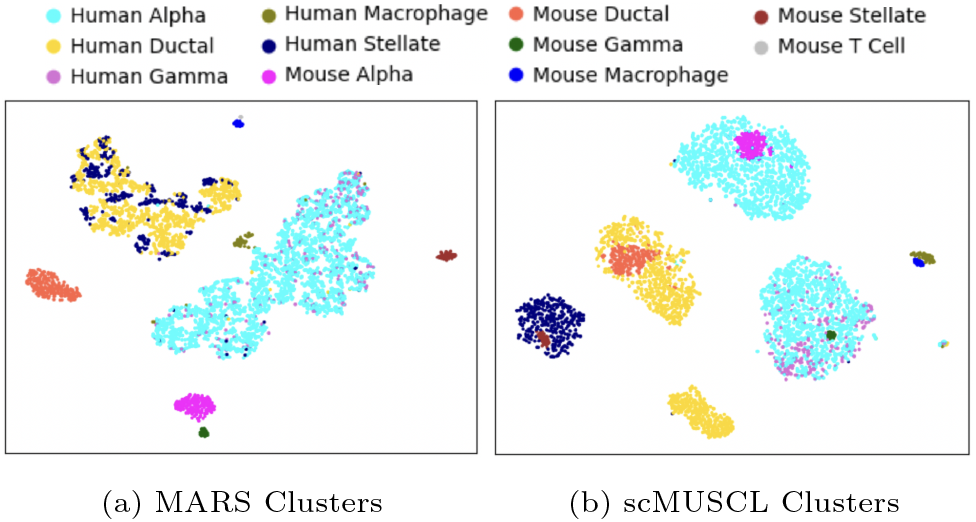
TSNE plot of cells in a mouse and a human pancreas cells when a mouse tissue is used as the source dataset to cluster human cells. (a) shows the clusters generated by MARS and (b) shows scMUSCL’s clustering result. This illustrates how MARS fails in aligning human and mouse cell. For example, MARS creates a cluster for mouse alpha cells far away from human alpha cells, while scMUSCL successfully aligns these two clusters. More interestingly, we can see scMUSCL aligns rare cell-types, such as human and mouse macrophage cells, while MARS fails to do so.

Transfer learning aims to improve the performance of a machine learning method on a target domain by transferring relevant knowledge from different but related source domains (Zhuang *et al*., 2020). Many transfer learning methods are based on pre-training and fine-tuning, where a feature extractor is trained on one or multiple relevant but different domain(s) and is fined-tuned on the target domain. Theoretical analysis (Ben-David *et al*., 2010) shows that in order to minimize the error of the target learner, the distribution distance between source and target domains should be minimized, which is done by learning domain invariant representations for both source and target domains, mainly through domain adversarial training (Ganin *et al*., 2016; Zhou *et al*., 2021). In another line of research, Zhao *et al*. (2020) study contrastive learning (Chen *et al*., 2020b) and conclude that contrastive learning disentangles domain and class information, and can be employed as a strong pre-training method for transfer learning.

In this paper, we present scMUSCL, a novel multi-source transfer learning method for clustering scRNA-seq data. scMUSCL exploits multiple annotated source scRNA-seq datasets to learn different cell-type characteristics and transfers the extracted knowledge to find cell clusters in a target dataset. scMUSCL is able to automatically determine the number of clusters and it is composed of three stages. The first stage is the contrastive pre-training of the feature extractor which initializes a feature space such that similar cells are mapped closer to each other. The second stage, the cluster initialization, is where we initialize two sets of clusters; one for all source datasets combined and one for the target dataset. Fine-tuning is the final stage where scMUSCL performs cell-level and cluster-level alignment to address distribution discrepancies, and in turn, batch effect. In this stage scMUSCL learns domain invariant feature representations, which is required to perform knowledge transfer from the source domains to the target domain. During fine-tuning scMUSCL further exploits the source datasets to learn a clustering for the target dataset. Fine-tuning is guided by a loss function with two terms. The first term performs a cell-level alignment. It iteratively aligns cells from common cell-types across different source datasets, and learns compact clusters in the source and target domains. The second term performs an iterative cluster-level alignment and aligns source and target clusters of common cell-types. Extensive experiments on 20 real-world human and mouse tissues show scMUSCL’s superiority over state-of-the-art scRNA-seq clustering methods. The main contributions of this paper are as follows:

- We propose scMUSCL; a novel multi-source transfer learning method for clustering scRNA-seq data.
- We perform extensive experiments and show scMUSCL successfully transfers knowledge across different species, various platforms, and tissues. We use 20 real-world datasets in our experiments and show that scMUSCL outperforms the previous state-of-the-art scRNA-seq clustering methods by a large margin.
- Our extensive experiments show that scMUSCL benefits from multiple source datasets as its learning reference. They also show scMUSCL effectively estimates the correct number of clusters.

## Related Works

In this section we review recent machine-learning methods for cell clustering, which we roughly categorize into two groups: unsupervised methods and transfer-learning methods.

### Unsupervised Methods

Earlier works mostly focus on dimensionality reduction, i.e. how to map high dimensional cell profiles into a much lower dimensional space for further downstream tasks such as clustering or visualization, (Wang *et al*., 2018; Lopez *et al*., 2018; Stuart *et al*., 2019; Tian *et al*., 2019; Ciortan and Defrance, 2021). One example is SIMLR (Wang *et al*., 2018). SIMLR learns a similarity measure for scRNA-seq data to perform dimension reduction. It combines multiple kernels to learn a distance metric that fits the structure of the data the best. SIMLR also employs a rank constraint in the learned cell-to-cell similarity and graph diffusion to address the challenge of high levels of dropout events. SIMLR couples clustering with the learning of a cell-cell similarity matrix and a respective low-dimensional (latent) representation. SIMLR shows promising results and has been used widely in the literature. In another work, authors employ deep generative neural networks to propose scVI (Lopez *et al*., 2018). scVI uses stochastic optimization and a variational auto-encoder (Kingma and Welling, 2013) to aggregate information across similar cells and genes, and to approximate the distributions that underlie the observed expression values. In a more recent deep-neural-networks-based approach, authors propose scDeepCluster (Tian *et al*., 2019). scDeepCluster aims to jointly learn feature representations and cell clustering via explicit modelling of scRNA-seq data generation using a denoising autoencoder. Autoencoders are a special kind of neural networks which are suitable for learning a low-dimensional latent representation of high-dimensional data. Denoising autoencoders capture more robust latent representations by learning to predict the original input given the randomly corrupted input. scDeepCluster trains a denoising autoencoder using a zero-inflated negative binomial (ZINB) loss function. To simultaneously learn a clustering of cells as well as feature representations, scDeepCluster employs Kullback-Leibler (KL) divergence on the latent space as described in the ‘deep embedded clustering’ (DEC) algorithm (Xie *et al*., 2016). scziDesk (Chen *et al*., 2020a) is another clustering method that utilizes denoising autoencoders. scziDesk uses a denoising autoencoder to characterize scRNA-seq data while proposing a soft self-training K-means algorithm to cluster the cell population in the learned latent space. The self-training procedure aggregates similar cells to obtain a more cluster-friendly latent space.

Similar to autoencoders, contrastive learning (Chen *et al*., 2020b) is a self-supervised method for learning latent representations. Contrastive learning encourages augmentations (views) of the same input to have more similar representations compared to augmentations of different inputs. Contrastive-sc (Ciortan and Defrance, 2021) is based on contrastive learning, and it aims to learn cell’s latent features that facilitate clustering. In Contrastive-sc, a neural network first learns a representation for each cell through a representation training phase. The representation is then clustered with a general clustering algorithm, such as K-means or Leiden community detection. scNAME (Wan *et al*., 2023) is a successor of Contrastive-sc and it incorporates a mask estimation task to a neighbourhood contrastive learning framework for cell representation learning. Since scRNA-seq data is noisy, the mask estimation task helps to reveal uncorrupted data structure and denoise the scRNA-seq data. scNAME also introduces a neighbourhood contrastive paradigm with an offline memory bank, which achieves intra-cluster compactness, yet inter-cluster separation. scNAME shows strong performance and outperforms previous state-of-the-art approaches for single-cell clustering.

### Transfer Learning

All the methods we introduced so far are unsupervised, meaning they only exploit an unannotated scRNA-seq dataset to learn complex intrinsic cell relationships. scmap (Kiselev *et al*., 2018) and Moana (Wagner and Yanai, 2018) are two examples of methods that take advantage of the annotated scRNA-seq datasets as their reference. scmap (Kiselev *et al*., 2018) combines three metrics, Pearson correlation, cosine distance, and Spearman correlation, to quantify the closeness between a cell and the centroids of cell clusters. It projects cells in a target dataset to a lower dimensional space and annotates them based on their correlation with the average cell-type specific gene expression in the source data. Moana (Wagner and Yanai, 2018) is based on support vector machines and uses a linear kernel on PCA-transformed annotated source data to cluster cells in the target data. scAdapt (Zhou *et al*., 2021) is based on domain adaptation paradigm and it uses adversarial domain adaptation to learn domain-invariant features using annotated source and unannotated target datasets. Domain-invariant features allow one to learn a task (e.g., cell-type annotation) from a source domain and then transfer it to a related target domain. scAdapt is unable to learn from more than one source domain, and it assumes the source and the target datasets follow the same category distribution, meaning it is unable to detect and find unseen cell-types: cell-types that do not exist in the source data but are present in the target data. A more similar appraoch to our work is MARS (Brbić *et al*., 2020), which is also a multi-source transfer learning method for single-cell clustering that does not require to know the correct number of clusters in advance. MARS considers multiple annotated datasets as the source dataset and one unannotated dataset as the target dataset, and transfers knowledge extracted from the source dataset to cluster cells in the target dataset. The feature extractor in MARS is a stacked auto-encoder and is trained using a k-means-like loss function to map cells to clusters.

## Method

### Problem Definition

We are given |*S*| labeled source datasets 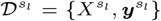 for 1 ≤ *l* ≤ |*S*| where 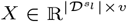 is a matrix containing 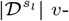 dimensional cells. *v* is the size of the input feature space, or here the number of genes. 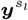 is a vector such that 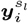 indicates the annotation (e.g., cell-type) for the *i*-th cell in the source dataset 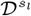. We are also given an unlabeled target dataset 𝒟^*t*^ = {*X*^*t*^} in which ***y***^***t***^ is not given. Our goal is to cluster cells in 𝒟^*t*^, such that each cluster represents either a seen or unseen cell-type. An unseen cell-type is a cell-type that exists in 𝒟^*t*^ but not in 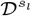 for 1 ≤ *l* ≤ |*S*|. Please note although this setting has similarities with the Universal Domain Adaptation (You *et al*., 2019) problem, our goal is to cluster the target dataset, not to classify it, which is the main objective in the Universal Domain Adaptation setting.

### scMUSCL

In this section we give an overview of scMUSCL, a transfer-learning-based clustering method to find cell clusters using scRNA-seq data. scMUSCL consists of a feature extractor *G*, which is a deep neural network that maps cells into a *d*-dimensional latent feature space where *d << v*. scMUSCL takes multiple annotated scRNA-seq experiments and one unannotated experiment as input, which we call the source datasets and the target dataset respectively. scMUSCL learns a clustering of the target dataset in three stages: constrastive pre-trainig, cluster initialization, and fine-tuning. In the first stage, we use the target and the source datasets to pre-train the feature extractor *G*. For pre-training we use InfoNCE contrastive loss (Oord *et al*., 2018). We randomly sample a minibatch of N cells and we create positive pairs of one cell derived from the minibatch and its augmentation, resulting in 2N cells. We obtain an augmentation of a cell by zeroing out non-zero gene expressions that are chosen randomly with some probability *p*. The InfoNCE loss for a positive pair of examples (*i, j*) is defined as:

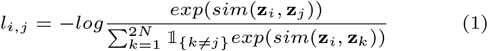

where 𝟙 denotes the indicator function. Here we define the similarity function as *sim*(**u, v**) = **u**^*T*^ **v***/*∥**u**∥∥**v**∥. We did not use a projection head and **z** is the direct output of our feature extractor *G*. Note that we omitted the temperature parameter from the InfoNCE loss formula since it’s equal to 1 in all our experiments. The final pre-training loss is computed across all positive pairs, both (i,j) and (j,i), in a mini-batch with size N. In the second stage of our method, we employ a pre-trained feature extractor, denoted as *G*, to calculate the latent representations of all cells. Utilizing these representations, we initialize two groups of cluster representatives: *C*^*s*^ for the combined source datasets, and *C*^*t*^ for the target dataset, both in ℝ^*K×d*^. Here, *K* represents the count of unique cell types across all source datasets. We define each row *k* in *C*^*s*^ as 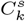 and express it as follows:

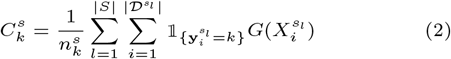

where 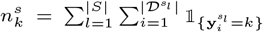 signifies the total number of cells in cell-type *k* across all source datasets. Following the initialization of *C*^*s*^, we duplicate its values to establish the initial values for the target cluster representatives, *C*^*t*^, such that *C*^*t*^ = *C*^*s*^ at this stage. This approach is predicated on the hypothesis that this pre-training yields a space where cells of the same type from source and target datasets are nearer to each other than cells of different types. In essence, if two cell groups, one from the source and the other from the target, share a cell-type, they should be proximal in this representation space. Consequently, it is logical to position two cluster representatives in this region—one for the source and another for the target. Nonetheless, we recognize that the target dataset may contain unseen cell-types, implying that not all target clusters will have a corresponding source cluster. These cells in the target dataset will initially align with their nearest target cluster representative and gradually shift the nearest representative towards themselves during the fine-tuning stage. It’s important to note that the number of target clusters is initialized to match the source clusters, though the actual number of target clusters is assumed to be fewer due to the amalgamation of multiple source datasets. This results in a surplus of initial target cluster representatives. If any representative in *C*^*t*^ fails to become the closest to any cells in the target dataset during fine-tuning, it remains unutilized. This mechanism allows scMUSCL to autonomously determine the requisite number of target clusters in the target dataset. In the last stage we fine-tune our feature extractor and optimize cluster representatives. We aim for 4 different goals in the fine-tuning stage:

1. To align cells with the same cell-type across different source datasets. Different source datasets may have cells with the same cell type, but their expression profiles may exhibit domain shift (e.g., batch effect). It is necessary to align these cells because otherwise the feature extractor might learn domain discriminative features, i.e., features that carry information about the domain rather than about the cell-type (Stojanov *et al*., 2021).
2. To learn compact target clusters. We seek to find compact clusters such that cells with the same cell-type are much closer to each other than cells with different cell-types.
3. To align source and target clusters of common cell-types. We assume the source datasets are highly relevant to the target dataset, which means they share some cell-types. We want to align these clusters with common cell-types in order to effectively transfer extracted knowledge from source datasets to the target dataset.
4. To learn well-separated clusters. We seek to find a representation space in which cells with different cell-types are located far apart.

Hence we propose a loss function with two terms. The first loss term, cell alignment loss, aims at the first two above goals. The second loss term, cluster alignment loss, aims at the latter two goals. In the following subsections we explain these two terms in detail.

#### Cell Alignment Loss

Cell alignment loss term minimizes the distance between the latent representation, 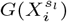, of *i*-th cell from source dataset 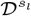 and its corresponding cluster representative. Source datasets are annotated, meaning we know the ground truth cell-type 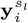 for each cell. Since source clusters are shared across all source datasets, minimizing the distance between a cell and its cluster representative leads to a cell-type-level alignment across source datasets. Moreover, it will help scMUSCL to learn compact clusters for source datasets. Cell alignment loss is illustrated in Figure 2a and is computed for each source dataset as defined below:

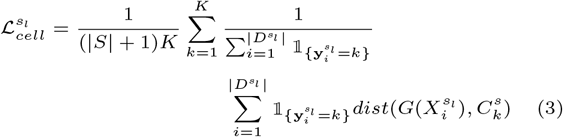

**Fig. 2.**
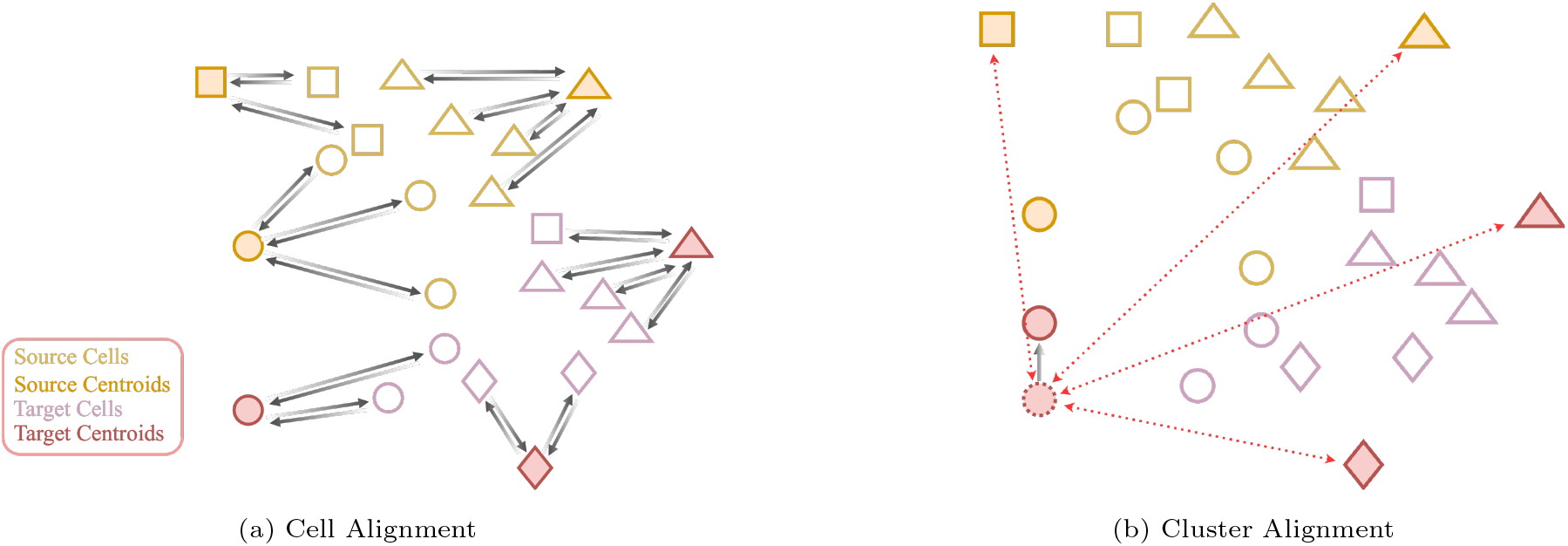
Illustration of scMUSCL’s fine-tuning loss function. Each shape represents a cell-type and filled shapes are cluster representatives. (a) Illustrates the Cell Alignment loss. Cell Alignment loss performs a cell-level alignment and aligns cells with the same cell-type across all source datasets and creates compact clusters in source and target domains. (b) Cluster Alignment loss performs a cluster-level alignment. It minimizes the distance between a cluster representative and its nearest cluster representative from the other domain while repelling all other cluster representatives.

We used Euclidean distance in our implementation for *dist*() but it could be any distance function. Note the coefficient 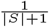 is to give similar weight to all the source and target datasets.

For the target dataset the loss function is similar, with the difference being that we minimize the distance between the latent features of an unannotated cell and its nearest cluster representative since we do not know the ground truth cell-type for it. We define the loss term for target cell alignment as below:

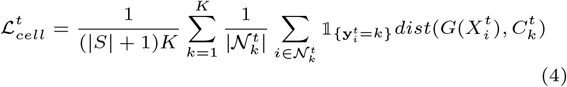

where 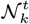 is the set of target cells that their nearest target cluster representative is *k*.

#### Cluster Alignment Loss

Enforcing cluster compactness is desirable, but it is also the recipe for collapse, as the feature extractor might minimize the loss by mapping all the cells and cluster representatives into one same vector. Therefore we need to ensure clusters are well-separated to avoid collapsing solutions. Moreover, we also need to introduce some kind of alignment between source and target clusters. scMUSCL learns two distinct sets of clusters; one for the union of source datasets, and one for the target dataset. These two cluster sets are optimized separately using cell alignment loss with no explicit and direct alignment across them. Therefore two clusters, one in the source set and one in the target set, might correspond to the same cell-type but yet be far apart because of the distribution difference between source and target datasets. Hence we need a strategy to align these clusters with common cell-types. The intuition behind this alignment is that it pushes the feature extractor to use the same set of genes to form the target cluster as the genes it used to form the source reference cluster of the same cell-type. This alignment will further translate the knowledge from the annotated source datasets to the unannotated target dataset. We propose a novel loss term, cluster alignment loss, based on entropy minimization. We assume that our pre-training and cluster representative initialization have already created a feature space with similar target and source clusters nearby. We want to encourage this closeness while we increase the inter-cluster distances among source and target clusters. We formalize this intuition by measuring the entropy of pair-wise cluster (representatives) similarities. We expect a target cluster to be similar to one source cluster (if that target cluster corresponds to a common cell-type) and dissimilar to the other source clusters and the other target clusters. A source cluster is expected to be dissimilar to the other source clusters. These assumptions imply a low entropy of the pairwise cluster similarities.

Cluster alignment loss is illustrated in Figure 2b. In order to compute it, we first define *F* ∈ ℝ^2*K×d*^ as 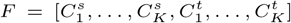. Having *F* defined, we define similarity matrix *P* such that for each value *p*_*i,j*_ in row *i* and column *j* we have:

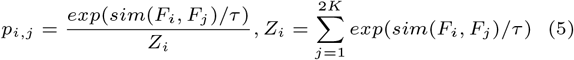

Here *sim*(*F*_*i*_, *F*_*j*_) is defined as below:

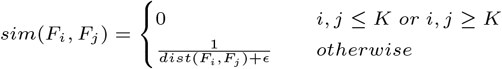

*ϵ* is a very small number added for numerical stability, and *τ* controls the distribution concentration degree (Hinton *et al*., 2015) which we set to 1 in our experiments. Now we can define our cluster alignment loss term as:

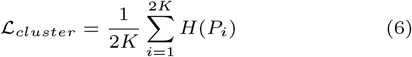

where *H*(*P*_*i*_) is the entropy of row *i* in *P*. Remember that K is the number of clusters in *C*^*s*^ and *C*^*t*^. By zeroing *p*_*i,j*_ for *i, j* ≤ *K or i, j > K* we are zeroing out the similarity among source clusters and the similarity among target clusters respectively. This way, we ensure that there is always a *j*^*′*^ such that *p*_*i,j*_ ≤ *p*_*i,j′*_ for (*i, j* ≤ *K or i, j > K*) and (*i* ≤ *K < j′ or j′* ≤ *K < i*). Therefore minimizing *H*(*Pi*) will increase the similarity with a *j′* with the highest *p*_*i,j′*_ and minimizes the similarity with all other cluster representatives.

### scMUSCL Loss Function

Having all the loss terms defined, the final loss function for the source and the target datasets is defined as follows:

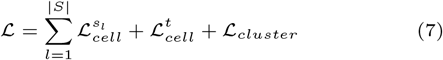

We use loss function ℒ to optimize the feature extractor parameters and cluster representatives in an alternating approach.

## Results

We designed a series of experiments to assess the performance of scMUSCL and compare it with five state-of-the-art methods. Among our baseline methods MARS (Brbić *et al*., 2020) is the most relevant to our work since it is also a transfer learning method that exploits multiple annotated source datasets. Two unsupervised methods based on contrastive learning, Contrastive-sc (Ciortan and Defrance, 2021) and scNAME (Wan *et al*., 2023) were included in the comparison, as this allows us to ask whether exploiting labeled source datasets in a transfer learning paradigm is beneficial. scDeepCluster (Tian *et al*., 2019) is also chosen as a baseline method since it is a deep-learning-based model that similar to scMUSCL learns feature representations and clustering simultaneously. The last baseline, SIMLR (Wang *et al*., 2018), is a method for learning cell representations particularly for clustering scRNA-seq data. We chose SIMLR as one of the baselines because it has been widely used in the literature. In all experiments of this section we used a feature extractor with two hidden layers with 1024 and 256 neurons, where the output layer has 100 neurons (*d* = 100). The reported performance results are the mean results of three independent experiments. The hyper-parameters of each experiment are shown in Supplementary Table 1. For SIMLR we used a python implementation available in *https://github.com/bowang87/SIMLR_PY*. For the remaining, we used their official implementations and default settings available in the corresponding Github page.

### Performance Metrics

We used Accuracy and Adjusted Rand Index (ARI) as our performance metrics. The Accuracy of clusterings is computed after running the Hungarian Maximum Matching Algorithm (Weber and Robinson, 2016) on the confusion matrix to find the best mapping between cell-types and cluster labels. Rand Index is a measure of the similarity between two data clusterings. It considers all pairs of samples and counts pairs that are assigned in the same or different clusters in the predicted and true clusterings. For example, true positive (TP) is defined as the number of sample pairs that belong to the same ground truth cluster and the same discovered (by the algorithm) cluster. Adjusted Rand Index is a form of the Rand Index (RI) which is adjusted for the chance grouping of elements. We used scikit-learn^1^ *metrics* API to compute these metrics.

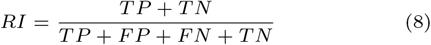

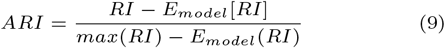

### Datasets

We used 20 real-world datasets to run our experiments. Human and mouse pancreas and kidney datasets were retrieved directly from SingleCellNet Github repository ^2^, except for Tabula Muris dataset which we downloaded directly from its official web page ^3^. We downloaded the human Peripheral Blood Mononuclear Cells (PBMC) scRNA-seq data from the SeuratData package (Ding *et al*., 2019). This data consists of seven batches from seven different sequencing platforms: cel-seq2, chrom-v3, indrop, smart-seq, chrom-v2, drop-seq, seq-well. We also used the dataset of 6 human endoderm-derived organs–lung, esophagus, liver, stomach, small intestine, and colon (Yu *et al*., 2021). List of these datasets and the details of our pre-processing is available in the Supplementary Materials.

### Cross-Platform Experiments

scRNA-seq data generated in different laboratories using different library preparation protocols and platforms often suffer from technical artifacts. Collectively, these batch effects can mask the biological information and limit the use of the available data. We first compared the performance of scMUSCL and baseline methods in their ability to transfer knowledge in the presence of batch effect across different sequencing technologies. For this, we used PBMC scRNA-seq data containing seven batches from seven different sequencing platforms: cel-seq2, chrom-v3, indrop, smart-seq, chrom-v2, drop-seq, seq-well. For each experiment, we assigned one dataset as the target dataset and the remaining six datasets served as the source datasets. When indrop platform was used as the target dataset, both scMUSCL and scNAME performed at a similar level. Importantly, in all remaining 6 experiments, scMUSCL achieved higher accuracy and ARI than all baseline methods (Figure 4a, Supplementary Table 5 and 6). Here we note even though both scMUSCL and MARS use transfer learning paradigm, the accuracy and ARI of scMUSCL was on average 0.35 and 0.43 higher respectively. This difference increases for chrom-v3 and chrom-v2, the two most common platforms in scRNA sequencing.

### Cross-Species Experiments

Having addressed that scMUSCL can effectively remove batch effects, we next asked whether scMUSCL can transfer knowledge from one species to another, hereon referred as cross-species. We carried out an experiment where we examined the difference in clustering efficiency measured by silhouette coefficient. For this, we used human pancreas tissue (Murano) as our unannotated target set and mapped it’s cells into the latent representation space where no training set was provided served as a source dataset. When one mouse pancreas dataset (Baron) was used for training, silhouette coefficient increased to 0.76 from baseline value of 0.03 (Figure 3). We next provided two mouse pancreas as training datasets (Baron and Tabula Muris), there was a modest increase in silhouette coefficient, showing that scMUSCL can effectively transfer knowledge between species. Comparable efficiency where one and two training datasets were used suggests that scMUSCL can be applied for clustering where limited number of training datasets are available.

**Fig. 3.**
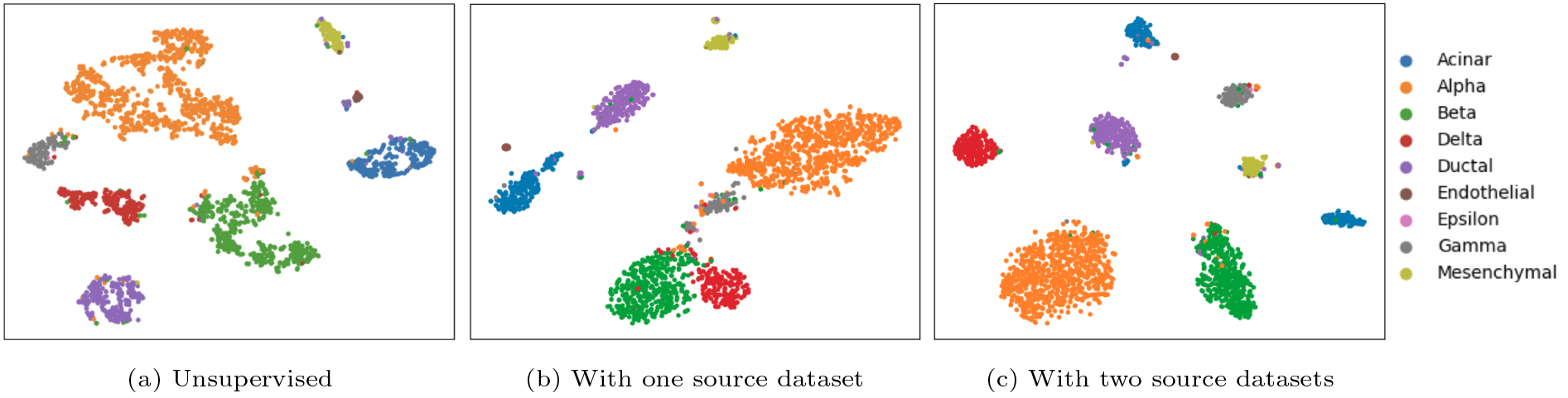
TSNE plots of cells in a human pancreas tissue (Murano). (a) Demonstrates cell clusters when we use no source dataset as the reference. We only use contrastive learning to learn the latent representations, silhouette score=0.03. In (b) we used one mouse pancreas tissue (Baron) as the source dataset, silhouette score=0.76, and in (c), we used two mouse pancreas source datasets (Baron and Tabula Muris), silhouette score=0.78.

Using accuracy and ARI as performance metrics, we next compared the ability of scMUSCL with baseline methods to transfer knowledge from mouse to human pancreas tissue. Here we used two mouse pancreas as training datasets (Baron and Tabula Muris) and three independent human pancreas target datasets. scMUSCL outperforms all of the baseline methods in terms of ARI and ACC in all three target dataset (Figure 4b and Supplementary Table 3). We in particular notice MARS’s poor performance on cross-species experiments. One explanation for this is that MARS fails to align source and target clusters in the presence of domain shift and therefore learns domain (instead of class) discriminative features (as depicted in Figure 1). This can significantly limit a transfer learning method’s performance and even lead to negative transfer. We also observe that unsupervised methods, SIMLR in particular, are showing comparable performance, especially in the experiment with Baron data as the target dataset. Despite this, unsupervised methods require the correct number of clusters in the target dataset, which is usually not available in practice.

**Fig. 4.**
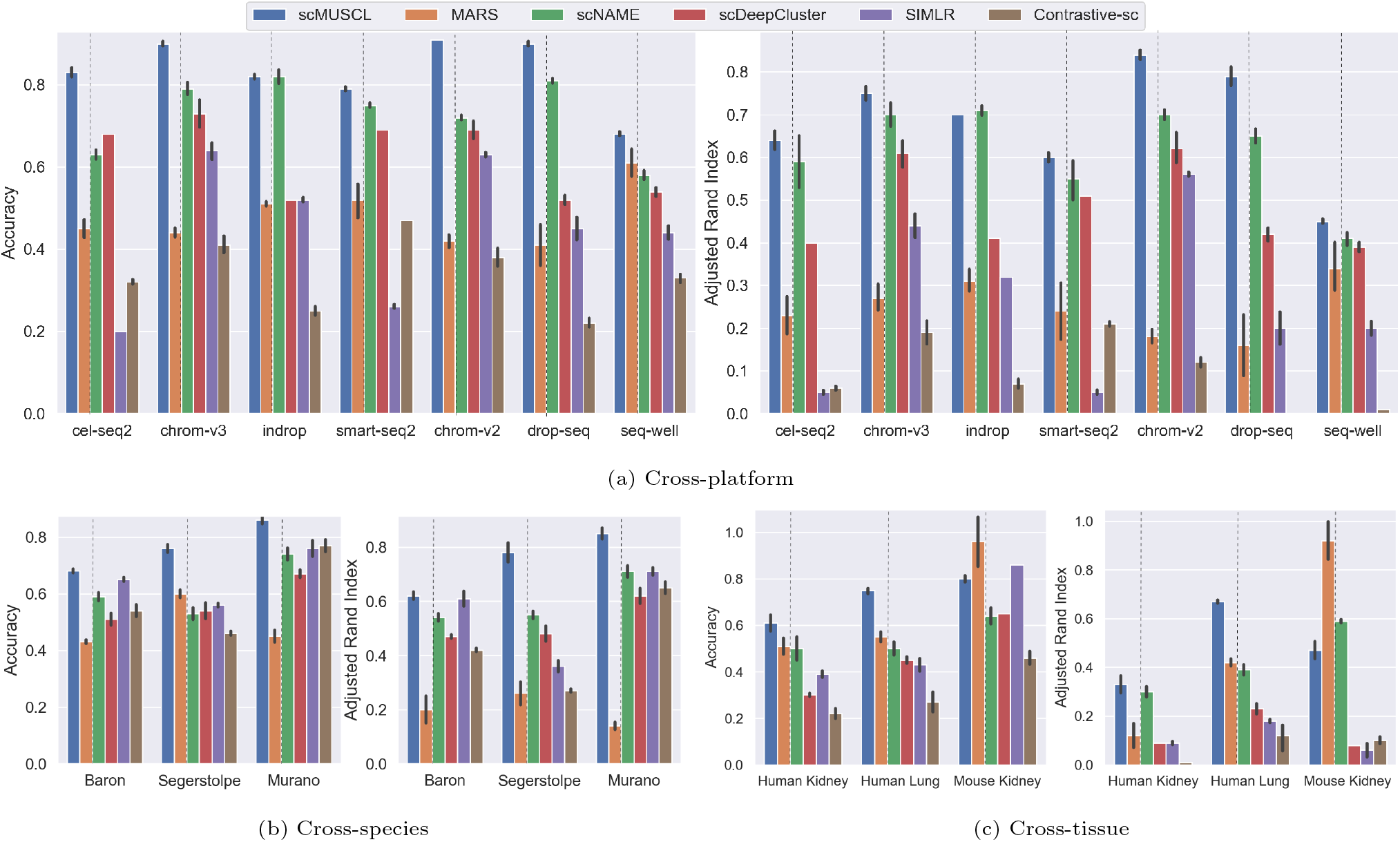
Accuracy and ARI of (a) cross-platform, (b) cross-species, and (c) cross-tissue experiments. The dotted line separates the transfer learning methods and unsupervised methods. (a) For cross-platform experiments we used 7 PBMC datasets. In each experiment we used one as the target dataset and exploited the rest as the source datasets. (b) In all cross-species experiments we used two mouse pancreas scRNA-seq data (Baron and Tabula Muris) as the source datasets, and one human pancreas as the target dataset. (c) For cross-tissue experiment we (i) exploited two human pancreas scRNA-seq datasets as the source dataset and one human kidney as the target dataset, (ii) one human esophagus and human short intestine as the source and a human lung as the target, and (iii) two mouse pancreas as the source and a mouse kidney as the target dataset.

### Cross-Tissue Experiments

We next analyzed the performance of scMUSCL on transferring between the tissue of the same species. When we used two mouse pancreas datasets (Baron, Tabula Muris) as source and one mouse kidney (Tabula Muris) as target datasets, both accuracy and ARI of MARS was higher scMUSCL. We note that MARS has been developed and evaluated on mouse cross-tissue experiments, we asked whether MARS can still perform better in an unforeseen datasets. For this, we used two human esophagus and short intestine tissues as training and human lung tissue as target dataset. scMUSCL had higher accuracy and ARI compared to MARS and other baseline methods, suggesting that the performance of MARS may be limited to the datasets used in developing the methodology. To consider an even more challenging case, we used two human pancreas datasets with approximately 7600 and 1900 cells as the source datasets, and a large human kidney dataset with more that 41000 cells as the target data. We see that even though scMUSCL has a relatively small source datasets available to learn from, it’s performance is still superior to all the baseline methods.

In order to evaluate whether scMUSCL and MARS can benefit from multiple source datasets, we performed five more cross-tissue experiments where we increased the number of source datasets by one in each experiment (Figure 5). To avoid bias in selecting the tissues to add into the training datasets, we selected a set of tissues, where diverse set of cell types, tissue-specific and shared, are present (Yu *et al*., 2021). We first used esophagus as the only source dataset, and we then add small intestine, colon, stomach, and liver to the source datasets and determined the effect of the number of source datasets in clustering of human lung tissue (Figure 5). We observed that scMUSCL’s performance improves with adding more source datasets, and its clustering accuracy increases from 0.62 with one source dataset to 0.91 with 5 source datasets. Although, MARS slightly outperforms scMUSCL when only one source dataset is used, it falls behind when more source datasets are added and is unable to even reach its initial accuracy. This again shows MARS inability in learning from multiple domains in the presence of batch effect, which leads to negative transfer.

**Fig. 5.**
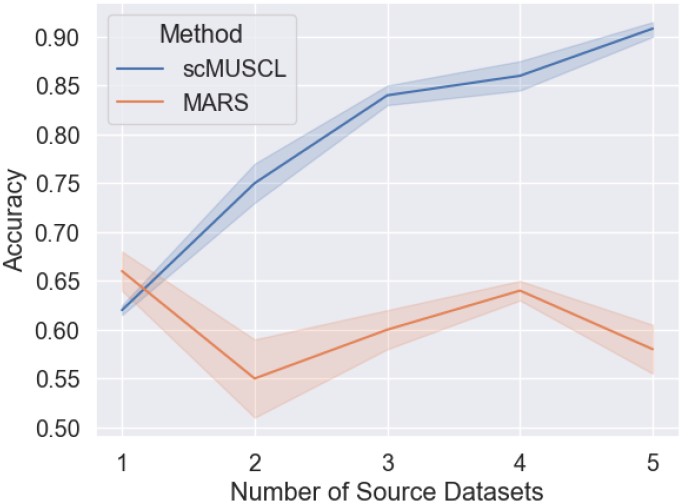
Accuracy of clustering on a human lung tissue with increasing number of endoderm-derived organs as the source datasets.

### Cluster Resolution

In contrast to other baseline methods, MARS and scMUSCL do not require user to provide the number of clusters in the target dataset in advance, and both are capable of automatically estimating the number of clusters in the target dataset during training procedure. Therefore, we compared the cluster resolution as a metric to evaluate MARS and scMUSCL in estimating the correct number of target clusters. Root Mean Square Error (RMSE) between the estimated number of target clusters and the ground truth number of clusters for all cross-species, cross-platform and cross-tissue experiments was lower when scMUSCL was used (Table 2), suggesting that scMUSCL is much more accurate in estimating the number of clusters in the target dataset compared with MARS.

**Table 1.** Results for three different versions of scMUSCL, removing one stage in each version.

| Ablation Studies: Cross-Species |  |  |  |  |  |  |
| --- | --- | --- | --- | --- | --- | --- |
| Pancreas Source | Mouse: Baron and Tabula Muris |  |  |  |  |  |
| Human Target | Baron |  | Seegerstolpe |  | Murano |  |
| Metric | ACC | ARI | ACC | ARI | ACC | ARI |
| scMUSCL w/o pre-training | $0.46 \pm 0.8$ | $0.18 \pm 0.12$ | $0.53 \pm 0.08$ | $0.14 \pm 0.16$ | $0.42 \pm 0.05$ | $0.10 \pm 0.14$ |
| scMUSCL w/o initialization | $0.49 \pm 0.03$ | $0.41 \pm 0.03$ | $0.42 \pm 0.02$ | $0.31 \pm 0.02$ | $0.52 \pm 0.00$ | $0.41 \pm 0.00$ |
| scMUSCL w/o fine-tuning | $0.36 \pm 0.00$ | $0.04 \pm 0.00$ | $0.52 \pm 0.01$ | $0.10 \pm 0.02$ | $0.43 \pm 0.00$ | $0.07 \pm 0.00$ |
| scMUSCL | **$0.68 \pm 0.01$** | **$0.62 \pm 0.02$** | **$0.76 \pm 0.02$** | **$0.78 \pm 0.05$** | **$0.86 \pm 0.02$** | **$0.85 \pm 0.03$** |

**Table 2.**
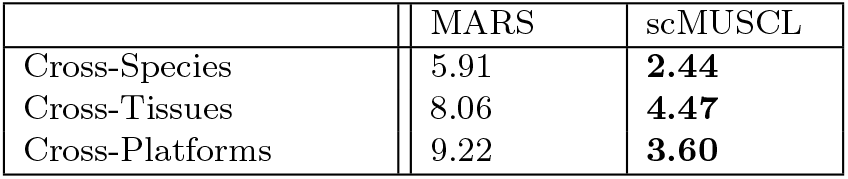
Root Squre Mean Error between the estimated and the ground truth number of target clusters.

|  | MARS | scMUSCL |
| --- | --- | --- |
| Cross-Species | 5.91 | **2.44** |
| Cross-Tissues | 8.06 | **4.47** |
| Cross-Platforms | 9.22 | **3.60** |

### Ablation Studies

We created three different versions of scMUSCL to evaluate the effectiveness of each of its three stages: (1) scMUSCL w/o pre-training, in which we did no contrastive pre-training, (2) scMUSCL w/o initialization, in which we used k-means to initialize target clusters, and (3) scMUSCL w/o fine-tuning, in which we did no fine tuning. As we can see in Table 1, scMUSCL benefits from each of these three stages. Moreover, we plot ACC and ARI of the target dataset and the source datasets during training in Figure 6. For this plot we used a cross-species experiment in which two mouse pancreas tissues from Baron and Tabula Muris are used as the source dataset and Murano human pancreas tissue is used and the target. We see in Figure 6b that our model starts from a high source accuracy, which is because contrastive pre-training has already learned some characteristics of the cells. However, source datasets are not aligned yet. The increasing trend in source accuracy indicates that our fine-tuning stage is aligning source clusters. Looking at Figure 6a we can see the target accuracy rapidly increases at the first few epochs. We believe this is because our model quickly learns to transfer the extracted knowledge from the source datasets to the target dataset. The target accuracy continues increasing as our model learns to align the source and target datasets.

**Fig. 6.**
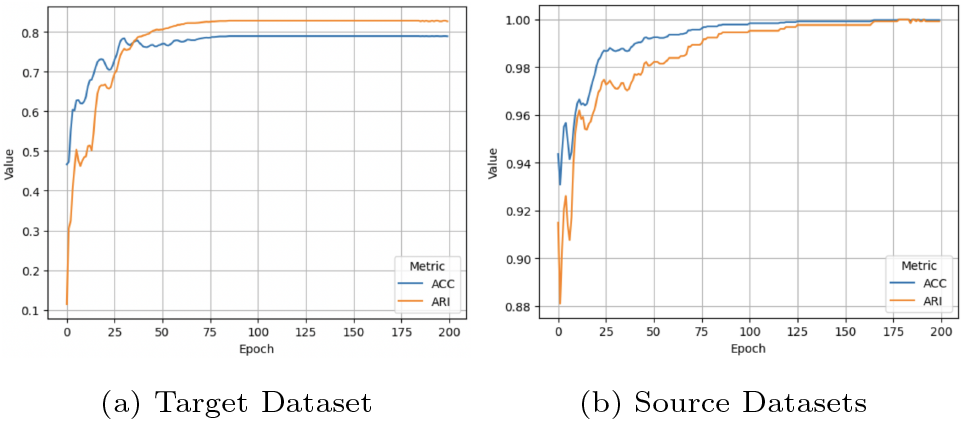
ARI and ACC of (a) target dataset and (b) source datasets in a cross-species experiment. Here we used two mouse pancreas datasets from Baron and Tabula Muris as source datasets and human pancreas dataset by Murano as the target dataset.

## Conclusion

In this paper, we investigated the problem of scRNA-seq data clustering and pointed out that the majority of current scRNA-seq clustering methods ignore the wealth of available annotated scRNA-seq datasets. We proposed scMUSCL, a multi-source transfer learning method that exploits multiple annotated scRNA-seq datasets as its reference to find clusters of cells in a target unannotated scRNA-seq dataset. We showed that scMUSCL does not require to know the number of target clusters beforehand. scMUSCL’s ability to estimate the correct number of clusters eliminates the need of additional tools. Considering that several laboratories are now adopting to scRNAseq technology, scRNAseq data obtained by different operators and platforms, therefore with batch effects will continue to raise. The use of scMUSCL can remedy this problem, as it can incorporate scRNAseq data coming from different platforms and can effectively remove batch effects. In parallel to this, we also showed that scMUSCL performance increases as we add relevant annotated scRNA-seq data to its source datasets. We designed and conducted extensive experiments and showed that scMUSCL can transfer knowledge across platforms, species and tissues and outperforms all the state-of-the-art scRNA-seq clustering methods by a large margin. scMUSCL effectively aligns clusters of cell types shared between the different source datasets and shared between the source datasets and the target dataset (if exist). We believe that this alignment is the main reason behind outperforming MARS (Brbić *et al*., 2020)–a state-of-the-art transfer learning-based method for scRNAseq clustering.

This paper suggests various directions for future work. scMUSCL implicitly gives the same weights to all the source datasets. While this is a reasonable default, depending on the biological question, different source datasets may have different relevance for the target dataset. scMUSCL can be adopted to weigh the source datasets by relevance by using their similarity to the target dataset. Furthermore, although scMUSCL can transfer from multiple source datasets, it is still limited to one target dataset. Extending scMUSCL to multiple target datasets promises to help it to better model the cell-cell relationships.

## Acknowledgements

We would like to show our gratitude for the help we received from Shuman Peng and Maedeh Abedi during this project.

## Footnotes

1 https://scikit-learn.org/

2 https://github.com/pcahan1/singleCellNet

3 https://tabula-muris.ds.czbiohub.org

## References

Ben-David, S. et al. (2010). A theory of learning from different domains. Machine learning, 79(1), 151–175.

Brbić, M. et al. (2020). Mars: discovering novel cell types across heterogeneous single-cell experiments. Nature methods, 17(12), 1200–1206.

Chen, L. et al. (2020a). Deep soft k-means clustering with self-training for single-cell rna sequence data. NAR genomics and bioinformatics, 2(2), qaa039.

Chen, T. et al. (2020b). A simple framework for contrastive learning of visual representations. In International conference on machine learning, pages 1597–1607. PMLR.

Ciortan, M. and Defrance, M. (2021). Contrastive self-supervised clustering of scrna-seq data. BMC bioinformatics, 22(1), 1–27.

Clarke, Z. A. et al. (2021). Tutorial: guidelines for annotating single-cell transcriptomic maps using automated and manual methods. Nature protocols, 16(6), 2749–2764.

Ding, J. et al. (2019). Systematic comparative analysis of single cell rna-sequencing methods. BioRxiv, page 632216.

Eberwine, J. et al. (2014). The promise of single-cell sequencing. Nature methods, 11(1), 25–27.

Ganin, Y. et al. (2016). Domain-adversarial training of neural networks. The journal of machine learning research, 17(1), 2096–2030.

Grabski, I. N. et al. (2023). Significance analysis for clustering with single-cell rna-sequencing data. Nature Methods, 20(8), 1196–1202.

Hinton, G. et al. (2015). Distilling the knowledge in a neural network. arXiv preprint arXiv:1503.02531.

Kingma, D. P. and Welling, M. (2013). Auto-encoding variational bayes. arXiv preprint arXiv:1312.6114.

Kiselev, V. Y. et al. (2018). scmap: projection of single-cell rna-seq data across data sets. Nature methods, 15(5), 359–362.

Lopez, R. et al. (2018). Deep generative modeling for single-cell transcriptomics. Nature methods, 15(12), 1053–1058.

Luecken, M. D. and Theis, F. J. (2019). Current best practices in single-cell rna-seq analysis: a tutorial. Molecular systems biology, 15(6), e8746.

Oord, A. v. d. et al. (2018). Representation learning with contrastive predictive coding. arXiv preprint arXiv:1807.03748.

Stojanov, P. et al. (2021). Domain adaptation with invariant representation learning: What transformations to learn? Advances in Neural Information Processing Systems, 34, 24791–24803.

Stuart, T. et al. (2019). Comprehensive integration of single-cell data. Cell, 177(7), 1888–1902.

Tian, T. et al. (2019). Clustering single-cell rna-seq data with a model-based deep learning approach. Nature Machine Intelligence, 1(4), 191–198.

Wagner, F. and Yanai, I. (2018). Moana: a robust and scalable cell type classification framework for single-cell rna-seq data. BioRxiv, page 456129.

Wan, H. et al. (2023). scname: neighborhood contrastive clustering with ancillary mask estimation for scrna-seq data. Bioinformatics, 38(6), 1575–1583.

Wang, B. et al. (2018). Simlr: A tool for large-scale genomic analyses by multi-kernel learning. Proteomics, 18(2), 1700232.

Wang, T. et al. (2019). Comparative analysis of differential gene expression analysis tools for single-cell rna sequencing data. BMC bioinformatics, 20(1), 1–16.

Weber, L. M. and Robinson, M. D. (2016). Comparison of clustering methods for high-dimensional single-cell flow and mass cytometry data. Cytometry Part A, 89(12), 1084–1096.

Xie, J. et al. (2016). Unsupervised deep embedding for clustering analysis. In International conference on machine learning, pages 478–487. PMLR.

You, K. et al. (2019). Universal domain adaptation. In Proceedings of the IEEE/CVF conference on computer vision and pattern recognition, pages 2720–2729.

Yu, Q. et al. (2021). Charting human development using a multi-endodermal organ atlas and organoid models. Cell, 184(12), 3281–3298.

Zhang, S. et al. (2023). Review of single-cell rna-seq data clustering for cell-type identification and characterization. RNA, 29(5), 517–530.

Zhao, N. et al. (2020). What makes instance discrimination good for transfer learning? arXiv preprint arXiv:2006.06606.

Zhou, X. et al. (2021). scadapt: Virtual adversarial domain adaptation network for single cell rna-seq data classification across platforms and species. bioRxiv.

Zhuang, F. et al. (2020). A comprehensive survey on transfer learning. Proceedings of the IEEE, 109(1), 43–76.

