## Supplementary Material for "scMUSCL: Multi-Source Transfer Learning for Clustering scRNA-seq Data"

### scMUSCL: Supplementary Materials

#### 1 Datasets and Pre-processing

We used 20 datasets to run our experiments. These datasets are listed in table 2. Please note PBMC dataset by Ding is actually 7 different batches sequenced by 7 different technologies. For each experiment we only kept the genes that are shared among source and target datasets. For Tabula Muris pancreas tissue we removed pancreatic A, D, and PP cell. We also removed 'immune other' from Baron mouse pancreas dataset and renamed 'activated-stellate', and 'quiescent-stellate' to 'stellate' in Baron mouse and human pancreas dataset. We further removed 'unclassified', 'co-expression', 'unclassified endocrine', and 'MHC class II' from Segerstolpe human pancreas dataset and renamed 'PSC' to stellate. In Park human kidney dataset we removed 'ascending loop of Henle', 'novel1', and 'novel2' cells and renamed 'proximal tubule', and 'distal convoluted tubule' to 'tubule'. We also renamed 'collecting duct intercalated cell', 'collecting duct principal cell', and 'collecting duct transitional cell' to 'collecting duct' and renamed 'endothelial, vascular, and descending loop of Henle' to endothelial. In Tabula Muris pancreas heart tissue we renamed 'smooth muscle cell', and 'cardiac muscle cell' to 'muscle'. Lastly, in six endodermal datasets by Yu et. a., we remove undefined cells, renamed 'T cell/NK cell 1' and 'T cell/NK cell 2' into 'T Cell', renamed 'Mesenchyme subtype 1' to 5 and 'Proliferative mesenchyme' to Mesenchyme, renamed 'PNS glia' and 'PNS neuron' to PNS, and renamed 'Distal lung epithelium', 'Gastrointestinal epithelium', and 'Intestinal epithelium' to Epithelium. We used ScanPy [1] python package to log normalize all datasets with the scale factor of 10000.

#### 2 Hyper-parameter Tuning

Chosen values for hyper-parameters of our method are shown in table 1. We did a grid search over a limited number of options for hyper-parameters in each category of experiments (cross-specie, cross-tissue, and cross-platform). For pre-training and fine-tuning learning rate we searched among [0.005, 0.001, 0.0005]. For pre-training and fine-tuning epoch number we searched among [60, 80] and [200, 300] respectively and we did no early stopping. For our feature extractor we tried two options. One with a hidden layer of 1024 neurons and one with two hidden layers with 1024 and 256 neurons. For the size of extracted latent representations we considered three options [80, 100, 128]. We also tried using a learning rate scheduler with gamma=0.1 and step size of 50.

#### 3 Results

Tables 3, 4, 6, 5 further show the results of our experiments. Each number is a mean value of three different runs. We also report further metrics for these experiments in Tables 7, 8, and 9.

[1] https://scanpy.readthedocs.io/en/stable/

| Experiment | Pre-training Epochs | Pre-training LR | Training Epochs | Training LR | LR Scheduler |
|---|---|---|---|---|---|
| Cross-Species | 60 | 0.0005 | 200 | 0.0005 | × |
| Cross-Tissue | 40 | 0.0005 | 200 | 0.005 | ✓ |
| Cross-Platform | 40 | 0.0005 | 300 | 0.0005 | × |

Table 1: Selected Hyper-parameter for our experiments.

| Specie | Tissue | Source |
|---|---|---|
| Mouse | Pancreas | Baron |
| Mouse | Pancreas | Tabula Muris |
| Human | Pancreas | Baron |
| Human | Pancreas | Segerstolpe |
| Human | Pancreas | Murano |
| Human | Kidney | Park |
| Mouse | Kidney | Tabula Muris |
| Human | PBMC | Ding |
| Human | Lung | Yu et. al. |
| Human | Esophagus | Yu et. al. |
| Human | Liver | Yu et. al. |
| Human | Stomach | Yu et. al. |
| Human | Small Intestine | Yu et. al. |
| Human | Colon | Yu et. al. |

Table 2: List of all 20 scRNA-seq datasets we used in our empirical study. Human PBMC dataset by Ding is actually 7 different batches sequenced by 7 different technologies.

| Cross-Species | | | | | | |
|---|---|---|---|---|---|---|
| Pancreas Source | Mouse: Baron and Tabula Muris | | | | | |
| Human Target | Baron | | Segerstolpe | | Murano | |
| Metric | ACC | ARI | ACC | ARI | ACC | ARI |
| scNAME | .0.59 ± 0.02 | 0.54 ± 0.02 | 0.53 ± 0.03 | 0.55 ± 0.02 | 0.74 ± 0.03 | 0.71 ± 0.03 |
| MARS | 0.43 ± 0.01 | 0.20 ± 0.07 | 0.60 ± 0.02 | 0.26 ± 0.06 | 0.45 ± 0.03 | 0.14 ± 0.02 |
| scDeepCluster | 0.51 ± 0.03 | 0.47 ± 0.01 | 0.54 ± 0.04 | 0.48 ± 0.04 | 0.67 ± 0.02 | 0.62 ± 0.04 |
| SIMLR | 0.65 ± 0.01 | **0.61 ± 0.04** | 0.56 ± 0.01 | 0.36 ± 0.03 | 0.76 ± 0.04 | 0.71 ± 0.02 |
| Contrastive-sc | 0.54 ± 0.03 | 0.42 ± 0.01 | 0.46 ± 0.01 | 0.27 ± 0.01 | 0.77 ± 0.03 | 0.65 ± 0.03 |
| scMUSCL | **0.68 ± 0.01** | **0.62 ± 0.02** | **0.76 ± 0.02** | **0.78 ± 0.05** | **0.86 ± 0.02** | **0.85 ± 0.03** |

Table 3: Results of cross-species experiment when using two mouse pancreas tissues as the source datasets and three independent human pancreas tissue as the target dataset. Reported numbers are mean and standard deviation over three runs.

| Cross-Tissues | | | | | | |
|---|---|---|---|---|---|---|
| Source | Human Pancreas: Baron, Segerstolpe | | Human Yu: Esophagus, short intestine | | Mouse Pancreas: Baron, TM | |
| Target | Kidney Park Human | | Yu Lung Human | | Kidney Tabula Muris Mouse | |
| Metric | ACC | ARI | ACC | ARI | ACC | ARI |
| scNAME | 0.50 ± 0.07 | 0.30 ± 0.03 | 0.50 ± 0.04 | 0.39 ± 0.03 | 0.64 ± 0.05 | 0.59 ± 0.01 |
| MARS | 0.51 ± 0.05 | 0.12 ± 0.07 | 0.55 ± 0.03 | 0.42 ± 0.02 | **0.96 ± 0.15** | **0.92 ± 0.11** |
| scDeepCluster | 0.30 ± 0.01 | 0.09 ± 0.00 | 0.45 ± 0.02 | 0.23 ± 0.03 | 0.65 ± 0.00 | 0.08 ± 0.00 |
| SIMLR | 0.22 ± 0.03 | 0.01 ± 0.00 | 0.27 ± 0.06 | 0.12 ± 0.09 | 0.46 ± 0.04 | 0.10 ± 0.02 |
| Contrastive-sc | 0.39 ± 0.02 | 0.09 ± 0.01 | 0.43 ± 0.04 | 0.18 ± 0.01 | 0.86 ± 0.00 | 0.06 ± 0.04 |
| scMUSCL | **0.61 ± 0.05** | **0.33 ± 0.05** | **0.75 ± 0.02** | **0.67 ± 0.01** | 0.80 ± 0.02 | 0.47 ± 0.05 |

Table 4: Results of three cross-tissue experiments. In the first experiment, we used two human pancreas tissues as source datasets and transfer the extracted knowledge to find clusters in a human kidney dataset. In the second experiment we learned from human esophagus and small intestine tissues and used a human lung tissue as the target dataset. In the third experiment we learned from two mouse pancreas datasets to cluster a human kidney dataset. Reported numbers are mean and standard deviation over three runs.

| Cross-Platform | | | | | | |
|---|---|---|---|---|---|---|
| Target | chrom-v2 | | drop-seq | | seq-well | |
| Metric | ACC | ARI | ACC | ARI | ACC | ARI |
| scNAME | $0.72 \pm 0.01$ | $0.70 \pm 0.02$ | $0.81 \pm 0.01$ | $0.65 \pm 0.03$ | $0.58 \pm 0.02$ | $0.41 \pm 0.03$ |
| MARS | $0.42 \pm 0.03$ | $0.18 \pm 0.03$ | $0.41 \pm 0.09$ | $0.16 \pm 0.13$ | $0.61 \pm 0.06$ | $0.34 \pm 0.11$ |
| scDeepCluster | $0.69 \pm 0.04$ | $0.62 \pm 0.07$ | $0.52 \pm 0.02$ | $0.42 \pm 0.03$ | $0.54 \pm 0.02$ | $0.39 \pm 0.02$ |
| SIMLR | $0.63 \pm 0.01$ | $0.56 \pm 0.01$ | $0.45 \pm 0.05$ | $0.20 \pm 0.07$ | $0.44 \pm 0.03$ | $0.20 \pm 0.03$ |
| Contrastive-sc | $0.38 \pm 0.04$ | $0.12 \pm 0.02$ | $0.22 \pm 0.02$ | $0.00 \pm 0.00$ | $0.33 \pm 0.02$ | $0.01 \pm 0.00$ |
| scMUSCL | $\mathbf{0.91 \pm 0.00}$ | $\mathbf{0.84 \pm 0.02}$ | $\mathbf{0.90 \pm 0.01}$ | $\mathbf{0.79 \pm 0.04}$ | $\mathbf{0.68 \pm 0.01}$ | $\mathbf{0.45 \pm 0.01}$ |

Table 5: Results of cross-platform experiments on a PBMC dataset. This data consists of seven batches from seven different sequencing platforms. Here we use one batch as the target datasets and the remaining 6 batches as the source dataset. Reported numbers are mean and standard deviation over three runs.

| Cross-Platform | | | | | | | | |
|---|---|---|---|---|---|---|---|---|
| Target | cel-seq2 | | chrom-v3 | | indrop | | smart-seq2 | |
| Metric | ACC | ARI | ACC | ARI | ACC | ARI | ACC | ARI |
| scName | $0.63 \pm 0.02$ | $0.59 \pm 0.11$ | $0.79 \pm 0.03$ | $0.70 \pm 0.05$ | $\mathbf{0.82 \pm 0.03}$ | $\mathbf{0.71 \pm 0.02}$ | $0.75 \pm 0.01$ | $0.55 \pm 0.09$ |
| MARS | $0.45 \pm 0.04$ | $0.23 \pm 0.08$ | $0.44 \pm 0.02$ | $0.27 \pm 0.06$ | $0.51 \pm 0.01$ | $0.31 \pm 0.05$ | $0.52 \pm 0.08$ | $0.24 \pm 0.12$ |
| scDeepCluster | $0.68 \pm 0.00$ | $0.40 \pm 0.00$ | $0.73 \pm 0.06$ | $0.61 \pm 0.06$ | $0.52 \pm 0.00$ | $0.41 \pm 0.00$ | $0.69 \pm 0.00$ | $0.51 \pm 0.00$ |
| SIMLR | $0.20 \pm 0.00$ | $0.05 \pm 0.01$ | $0.64 \pm 0.04$ | $0.44 \pm 0.05$ | $0.52 \pm 0.01$ | $0.32 \pm 0.00$ | $0.26 \pm 0.01$ | $0.05 \pm 0.01$ |
| Contrastive-sc | $0.32 \pm 0.01$ | $0.06 \pm 0.01$ | $0.41 \pm 0.04$ | $0.19 \pm 0.05$ | $0.25 \pm 0.02$ | $0.07 \pm 0.02$ | $0.47 \pm 0.00$ | $0.21 \pm 0.01$ |
| scMUSCL | $\mathbf{0.83 \pm 0.02}$ | $\mathbf{0.64 \pm 0.04}$ | $\mathbf{0.90 \pm 0.01}$ | $\mathbf{0.75 \pm 0.03}$ | $\mathbf{0.82 \pm 0.01}$ | $0.70 \pm 0.00$ | $\mathbf{0.79 \pm 0.01}$ | $\mathbf{0.60 \pm 0.02}$ |

Table 6: Results of cross-platform experiments on a PBMC dataset. This data consists of seven batches from seven different sequencing platforms. Here we use one batch as the target datasets and the remaining 6 batches as the source dataset. Reported numbers are mean and standard deviation over three runs.

| Cross-Species | | | | | |
|---|---|---|---|---|---|
| target | adj-mi | recall | nmi | precision | f1-score |
| Human Pancreas Baron | $0.748 \pm 0.021$ | $0.622 \pm 0.059$ | $0.749 \pm 0.021$ | $0.619 \pm 0.019$ | $0.566 \pm 0.020$ |
| Human Pancreas Segerstolpe | $0.832 \pm 0.037$ | $0.720 \pm 0.057$ | $0.834 \pm 0.037$ | $0.761 \pm 0.066$ | $0.722 \pm 0.059$ |
| Human Pancreas Murano | $0.825 \pm 0.022$ | $0.687 \pm 0.035$ | $0.827 \pm 0.021$ | $0.672 \pm 0.051$ | $0.672 \pm 0.041$ |

Table 7: Further metrics for cross-species experiments. Reported numbers are mean and standard deviation over three runs.

| Cross-Tissue | | | | | |
|---|---|---|---|---|---|
| target | adj-mi | recall | nmi | precision | f1-score |
| Human Kidney Park | $0.505 \pm 0.040$ | $0.456 \pm 0.055$ | $0.505 \pm 0.040$ | $0.442 \pm 0.026$ | $0.419 \pm 0.038$ |
| Human Lung Yu | $0.683 \pm 0.016$ | $0.656 \pm 0.021$ | $0.684 \pm 0.009$ | $0.659 \pm 0.019$ | $0.619 \pm 0.021$ |
| Mouse Kidney TM | $0.489 \pm 0.080$ | $0.807 \pm 0.056$ | $0.500 \pm 0.078$ | $0.980 \pm 0.028$ | $0.879 \pm 0.022$ |

Table 8: Further metrics for cross-tissue experiments. Reported numbers are mean and standard deviation over three runs.

| Cross-Platform | | | | | |
|---|---|---|---|---|---|
| target | adj-mi | recall | nmi | precision | f1-score |
| cel-seq2 | $0.718 \pm 0.016$ | $0.829 \pm 0.022$ | $0.725 \pm 0.016$ | $0.844 \pm 0.010$ | $0.827 \pm 0.010$ |
| chrom-v3 | $0.831 \pm 0.010$ | $0.921 \pm 0.004$ | $0.832 \pm 0.010$ | $0.889 \pm 0.001$ | $0.899 \pm 0.003$ |
| indrop | $0.735 \pm 0.004$ | $0.889 \pm 0.004$ | $0.736 \pm 0.004$ | $0.753 \pm 0.009$ | $0.781 \pm 0.010$ |
| smart-seq2 | $0.745 \pm 0.012$ | $0.867 \pm 0.010$ | $0.752 \pm 0.012$ | $0.865 \pm 0.008$ | $0.843 \pm 0.012$ |
| chrom-v2 | $0.851 \pm 0.009$ | $0.942 \pm 0.002$ | $0.852 \pm 0.009$ | $0.875 \pm 0.012$ | $0.897 \pm 0.007$ |
| drop-seq | $0.782 \pm 0.014$ | $0.900 \pm 0.011$ | $0.783 \pm 0.014$ | $0.820 \pm 0.011$ | $0.827 \pm 0.011$ |
| seq-well | $0.552 \pm 0.014$ | $0.817 \pm 0.010$ | $0.554 \pm 0.014$ | $0.739 \pm 0.010$ | $0.729 \pm 0.011$ |

Table 9: Further metrics for cross-platform experiments. Reported numbers are mean and standard deviation over three runs.
